# FECDO-Flexible and Efficient Coding for DNA Odyssey

**DOI:** 10.1101/2024.02.18.580107

**Authors:** Fajia Sun, Long Qian

## Abstract

DNA has been pursued as a compelling medium for digital data storage during the past decade. While large-scale data storage and random access have been achieved in artificial DNA, the synthesis cost keeps hindering DNA data storage from popularizing into daily life. In this study, we proposed a more efficient paradigm for digital data compressing to DNA, while excluding arbitrary sequence constraints. Both standalone neural networks and pre-trained language models were used to extract the intrinsic patterns of data, and generated probabilistic portrayal, which was then transformed into constraint-free nucleotide sequences with a hierarchical finite state machine. Utilizing these methods, a 12%-26% improvement of compression ratio was realized for various data, which directly translated to up to 26% reduction in DNA synthesis cost. Combined with the progress in DNA synthesis, our methods are expected to facilitate the realization of practical DNA data storage.

## Introduction

The exponentially growth of the global datasphere has been stimulating the demand of exploring novel storage media since the beginning of twenty-first century [1–4]. Inspired by the storage of biological information, DNA has been employed as functional carrier for digital data in recent years, which surpasses traditional storage media in density and durability [5–14]. However, the cost of storing large-scale data in DNA remains unaffordable despite the constant decrease in expenses of DNA synthesis and DNA sequencing, which hinders the popularization of DNA data storage [3, 15]. Specifically, the consumption of data storage in DNA is mainly derived from data writing (*i.e.*, DNA synthesis), which counts for more than 90% in total storage cost [16–17]. Furthermore, the cost of DNA synthesis is usually proportional to the number of nucleotides to be synthesized. Therefore, a vital aspect to reduce the cost of DNA data storage is to shorten the length of the nucleotide sequence required when storing specific data. Following this thought, two approach are supposed to be considered. The first one is data transcoding, namely, transforming data (normally in the form of binary stream) into nucleotide sequence. Various encoding methods have been explored during the past decade [5–15]. However, the natural bound of 2 bit per nucleotide confined the utility of this approach [16–17]. Another approach is data compression, namely, representing data in a more compact form. Previous studies of DNA data storage used compression methods adapted from information science, such as Huffman coding [6, 18], run-length coding [15, 19] or integrated algorisms like bzip2 [10, 20] (supplementary note 1). While these methods managed to condense digital data, the maximum possible extent of compression is not actually achieved [21–22]. This deficiency is especially detrimental for DNA data storage since the compression ratio is directly related to the cost.

Two issues are responsible for the inadequate compression of digital data into DNA. The first one is that data are considered as discrete and memoryless sequence in some compression methods (*e.g.*, Huffman coding) [18, 23]. However, digital information in the real world generally possesses correlation to the context, whose information entropy is overestimated [24]. Since information entropy denotes the extreme extent of data compression, as illustrated by Shannon’s first theorem [21], this overestimation leads to a longer sequence for storage (supplementary note 2). Second, nucleotide sequences possessing specific features are supposed to be excluded for data storage due to the difficulty during the synthesis and sequencing [5–6]. These features include homopolymer runs (*e.g.*, TTTTT), inappropriate secondary structure, heavily repeated subsequences, etc. Some studies attempted to avoid these sequences by designing special sequence screening algorithms [8, 14]. However, these algorithms resulted in a decrease of the storage density, since the sequence space available for encoding is reduced.

In this study, different from the traditional coding methods, we present FECDO (Flexible and Efficient Coding for DNA Odyssey), which is a more efficient and flexible compressive encoding paradigm for DNA data storage. The compression pipeline of FECDO is comprised of two main modules, a pattern-extraction module and a sequence-transformation module. The pattern-extraction module is based on neural networks, where we respectively use standalone neural networks and pre-trained language models to extract the feature of data to be storage. The sequence-transformation module is a hierarchical finite state machine, which excludes arbitrary sequence constrains with minimal overhead. To explicitly illustrate the superiority of this coding scheme, we compress a variety of digital data with FECDO and coding pipelines in previous studies of DNA data storage. As indicated by the results, FECDO enables persuasive degree of data compression, which outperforms both solus compression methods (*e.g.*, Huffman coding) and integrated compression methods (*e.g.*, bzip2). Specifically, compared with a pipeline comprised of bzip2 [10] and DNA fountain [8], FECDO realizes a 12%-26% reduction in compressed data size. Utilizing FECDO, we store a piece of text (10MB in size) into a minimal length of nucleotide sequence for synthesis, and achieve faithful information retrieval through nanopore sequencing. These results highlight the potential of FECDO in the field of DNA data storage, to effectively reduce the amount of DNA required to store information, thereby competently reducing the cost of DNA data storage.

## Results

### Overview of the FECDO pipeline

Digital data possess natural inherent correlation regardless of the specific format, which can be utilized for more efficient data compression. Intuitively, the masked part of the data can be predicted given careful inspection of a portion of the unmasked data. To exploit this correlation, the inner pattern of the data was first extracted using neural networks. Specifically, a probability model was built to characterize the contextual feature of a specific data sequence. Compression was then realized by sampling from the probability space of the model, resulting in conditional probabilities. These conditional probabilities were then mapped to a probability space represented by nucleotide sequences. This nucleotide sequence may contain a multitude of sequence constraints, which cannot be eliminated by merely defining mapping rules. Therefore, additional operations were performed to the sequence to exclude the constraints, which inevitably added redundancy due to the mapping to a higher-dimensional space and then contraction to the available space.

Figure 1 shows the workflow of compressing digital data to DNA sequence using FECDO. First, the digital data (text, images, videos, etc.) were converted into mathematical forms, that is, a tensor. Next, neural networks were used to build a probability model for the data, which was realized by training networks with the data tensor as inputs. After training, the characteristics of the data were stored in the weights of the networks. These weights were then recorded and used for prediction during both encoding and decoding process. As for encoding, the value of a specific location in the data sequence was obtained by feeding the upstream sequences to the trained network, and returning a conditional probability. This process was iteratively done until all the positions in data sequence were predicted, resulting in a string of conditional probabilities. To transform these conditional probabilities into nucleotide sequences, base-4 arithmetic coding was used over a probability space represented by four nucleotides. Finally, the resulting nucleotide sequences were processed by base shifter, a specially-designed machine, generating new sequence without constraints. This new sequence can be synthesized and then stored in DNA pools or live cells.

The decoding process required the retrieved nucleotide sequences and the stored weights of the neural networks. Specifically, the nucleotide sequences were first mapped to a particular region of the probability space. To restore a specific position in the data sequence, the upstream sequences were fed into the network with frozen weights, generating a conditional probability. By comparing the location of the probability region and the conditional probability, the correct value of the position was recognized and appended to the restored data sequence (Figure S1). This process was iteratively done for each position in data sequence until the whole sequence was restored.

**Figure 1.**
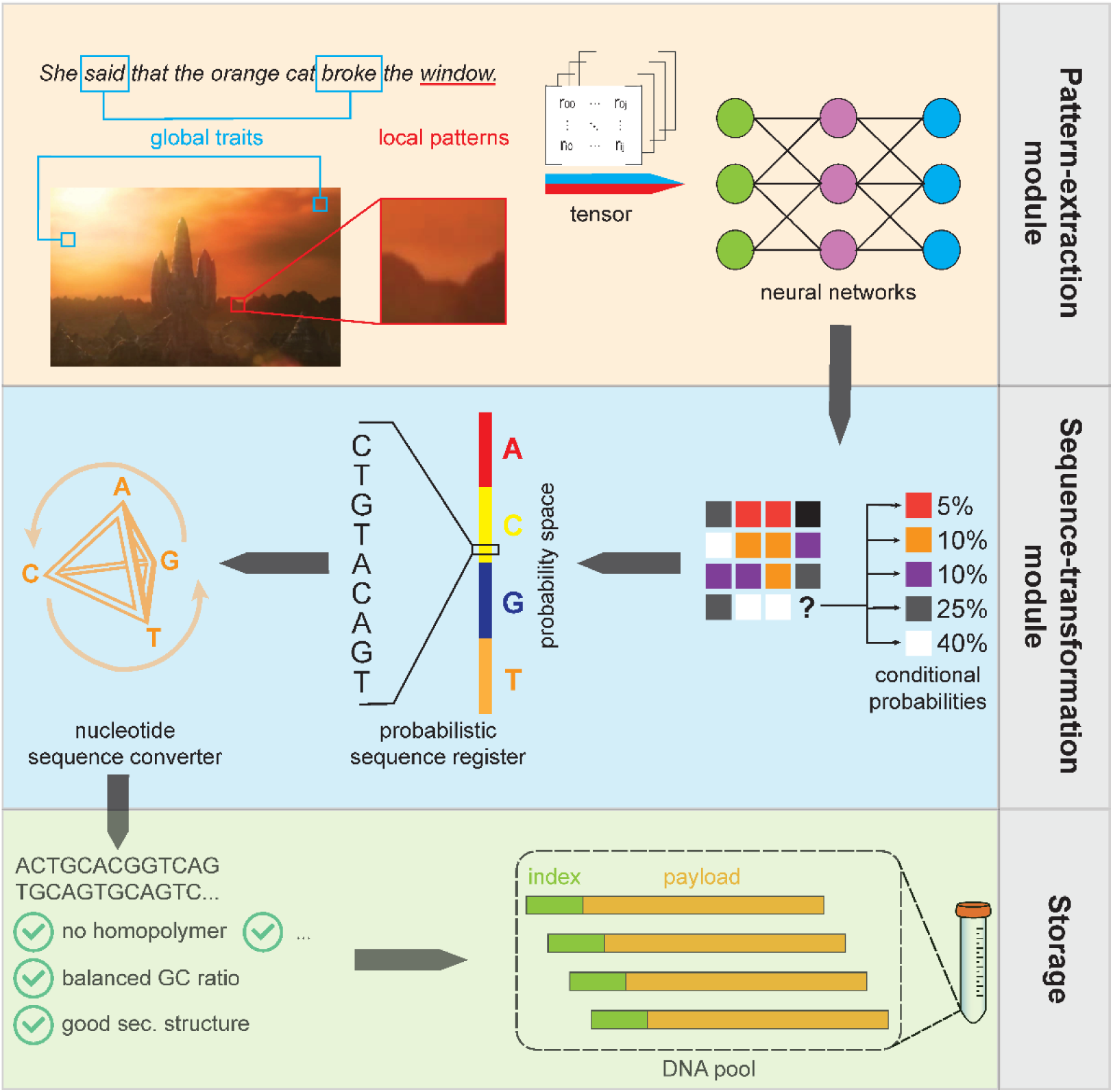
The workflow of FECDO. Digital data are first converted into tensor, and then fed into neural networks. After training, the characteristics of the data are stored in the parameters of the networks, which would generate conditional probabilities at of specific locations in data sequence. Next, these probabilities are transformed into nucleotide sequences, and the sequence constraints are then excluded to obtain constraint-free nucleotide sequences to be stored.

In the remaining sections, we used three text data, one grayscale image and one colored image to illustrate the coding efficiency of FECDO. The first text was a “clean” version of Wikipedia text [30] containing only lower-case English letters and spaces, which served as an exemplification of low-complexity English text (denoted as “wiki0”). The second text was the first 10,000,000 characters in The Intelligent Web Corpus [31], which was regarded as high-complexity English text (denoted as “web”). The third text was the first 10,000,000 characters in The Corpus do Portuguese [32], which was written in pure Portuguese and served as a representative of non-English text (denoted as “por”). The original size of each of these three texts was 10 MB. The shape of the grayscale image is 4032×3024, totaling 12,192,768 pixels. As each pixel corresponds to 8 bits (1 byte), the image size is 12.19 MB. The shape of the colored image is identical to that of grayscale image. However, each pixel corresponds to 24 bits (3 byte) in the colored image, resulting in an image size of 36.58 MB.

### Standalone neural networks for pattern extraction

We first devised a standalone network to model data sequences. Here “standalone” indicates that the network contains no prior knowledge of the data to be processed. In this case, the probability features of the sequence were completely reserved in the parameters of the network, and compression was simultaneously achieved through effectual sequence modelling. Since the parameters of the network were supposed to be stored for de-compression, the network must be light enough, while capable of effectively extract the pattern of data. These two considerations resulted in a trade-off, namely, studying a reasonably effective representation with fairly less parameters. In addition, as mentioned before, the model should adequately study both local and global features. Given these concerns, we constructed a model comprised of tandem Gated Recurrent Units (GRU) [25–26] and Transformer encoder [27–28] for sequence modelling (Figure 2a). The data sequence was split into short fragments and fed into GRU cells, whose outputs were concatenated and processed by Transformer encoder (Figure 2a). To further reduce the size of the parameters, we stored the parameters in float16 format rather than the default float32 [29]. In this case, the parameters were appended to float32 format before they can be computed with the tensor, and the computing result were truncated to obtain float16 format for storage (Figure 2b).

To verify the capability of this standalone network, we inputted the three representative texts into it. The pattern-extraction efficiency can be inspected by the “virtual” file size after compression using the network, which was calculated as

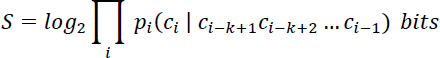

where *p* denoted conditional probability at the *i*-th position in the data sequence; *c_i_*denoted the correct character at the *i*-th position in the data sequence; *k* was a hyperparameter denoting the length of upstream sequences used for prediction.

Alternatively, *S* can also be directly obtained via the training loss

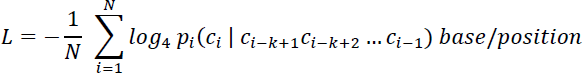

where *N* was the length of data sequence. Noting that this expression was essentially interchangeable to the expression of *S*. Figure 2c-e illustrate the “virtual” compressed size during the training process for the three texts. For comparison, the file sizes compressed by bzip2 are respectively indicated as black dash lines in each figure. While we adopted an optimized training setting where the learning rate was gradually reduced (Methods), the loss stagnated from around 50^th^ epoch. The compressed data sizes calculated based on this loss were 2.588 MB, 2.699 MB and 2.740 MB for the three texts, whereas the bzip2 compressed sizes were 2.660 MB, 2.808MB and 2.886 MB, respectively (Figure 2c-e). In other words, the compression efficiency of the network in Figure 2a is 2.71%, 3.88% and 5.05% higher than bzip2 in terms of representatives of low-complexity English text, high-complexity English text and non-English text, respectively (Figure 2i).

We also examined the capability of this network extracting the patterns in images. Specifically, we separately fed a grayscale image and a colored image into the network. Since the grayscale image was a two-dimensional array (with values ranging from 0 to 255), we simply flattened the array and then fed into the network in a manner similar to that of text. The colored image was a three-dimensional array, with the length of last panel being three (RGB). Therefore, we first split the array into three three-dimensional arrays, which were then flattened and concatenated and fed into the network (Figure 2f). For either image, the network successfully learned specific features, which was illustrated by the descending loss (Figure 2g&h). Noting that FECDO denotes a lossless data storage pipeline, the compression result can only be compared to other lossless methods, such as PNG (dashed line in Figure 2g&h). In contrast, the more commonly used JPEG is lossy, thus cannot serve as a baseline. The compression efficiency is 5.36% and 10.51% higher than PNG in terms of representatives of greyscale image and colored image, respectively (Figure 2i).

**Figure 2.**
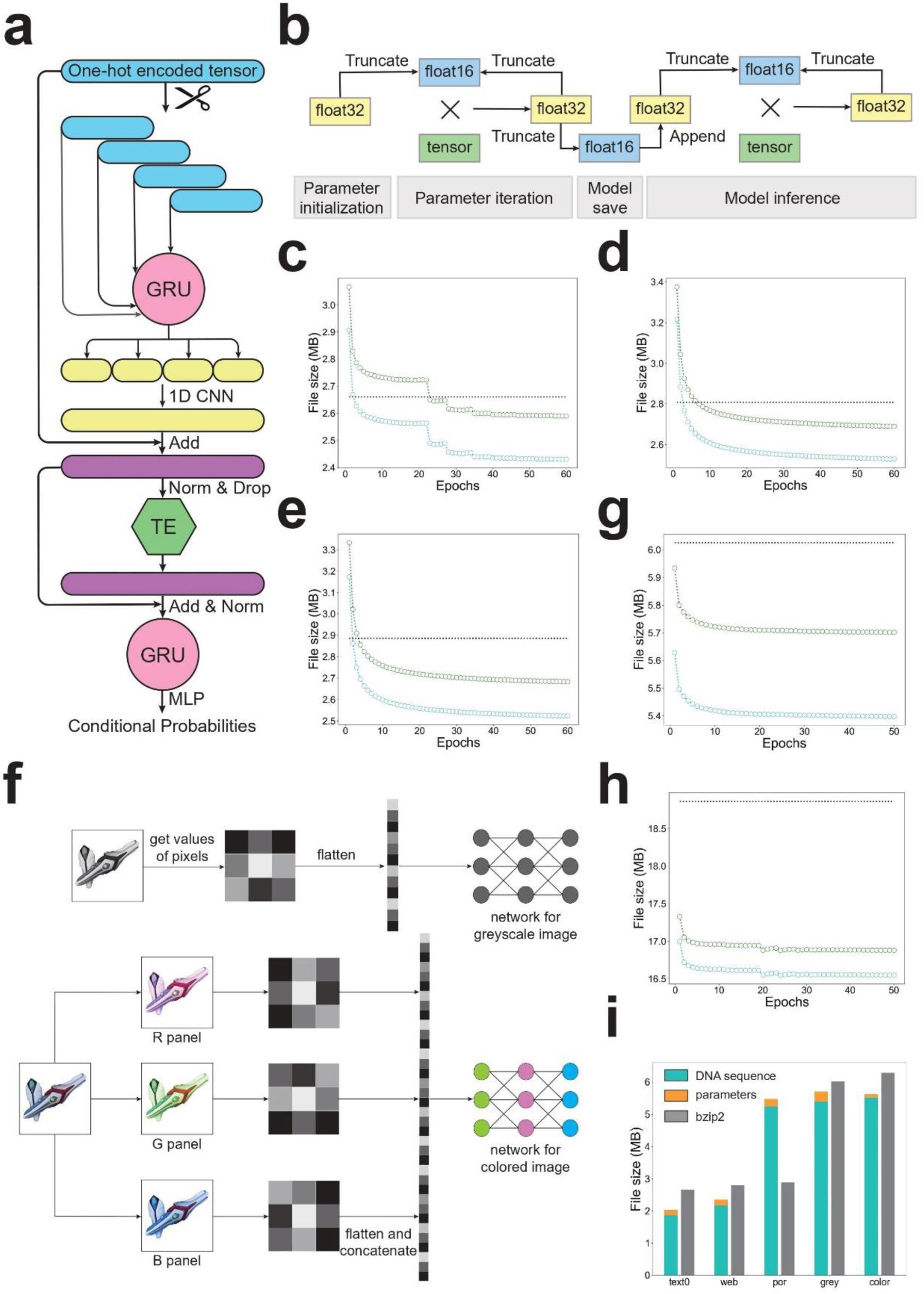
Pattern extraction using standalone network. (a) Structure of the network. TE: Transformer encoder. CNN: Convolutional neural network. Norm: Normalization. Drop: Dropout. Add: Residual connection. (b) The precision of parameters during training, storage and inference. (c)-(e) The “virtual” compressed size during the training process of (c) wiki0, (d) web and (e) por. In each panel, the cyan dots indicate the file size without the size of parameters, while the green dots indicate the file size with the size of parameters, and the black dash line indicates the file size compressed by bzip2. (f) The pre-processing of images (two-dimensional data). (g)-(h) The “virtual” compressed size during the training process of (g) greyscale image, (h) colored image. In each panel, the cyan dots indicate the file size without the size of parameters, while the green dots indicate the file size with the size of parameters, and the black dash line indicates the file size compressed by PNG. (i) The total compressed size of three texts and two images, in comparison with bzip2 and PNG. The values of last column (colored image) were divided by three to better illustrate other columns.

### Pattern extraction facilitated by pre-trained language models

Pattern extraction using the above network realized effective compression regardless of the content of the data. However, priori knowledge can be utilized for more efficient compression for specific data. For instance, the content of text can be better predicted if the rules of word formation were aware of. To fully exploit known knowledge for data compression, we further performed pattern extraction by prompting pre-trained language models (PLM). PLMs, like GPT [33] BERT [34] and Roberta [35], contain abundant information about natural language, since these models are trained with vast amount of text corpus. PLMs are usually applied to downstream tasks by fine-tuning, which is impractical for data compression due to the demands for storing large numbers (millions to billions) of model parameters. In addition, the tokenizers of these PLMs usually work at word or sub-word level, which resulted in a large number of cells required in the bottom layer due to numerous unique tokens (*e.g.*, more than 80,000 cells in the bottom layer of GPT2). The cost for training the network is therefore significantly increased, considering the computational complexity and the space required to store parameters.

To cope with these two difficulties, we here used a character-level PLM rather than the common word-level PLM, and we adopted a training method of “side network” or “prompt” [36]. Specifically, the PLM we used was Reformer [37], with an auxiliary network attached, forming a “composite network” (Figure 3a). This auxiliary network was comprised of two fully-connected (FC) layers. While a more delicate structure was devised for fine-tuning in a similar manner [38], we found the simple FC layers is sufficient for effectively extracting the patterns of input data. During the training process, the parameters of the PLM were frozen, and only the parameters of the auxiliary network were adapted. Given priori knowledge stored in the PLM, the compression efficiency is supposed to be further improved.

To confirm the above assumption, we used the composite network to extract the pattern in the three texts we previously used in the standalone network. The training setting was also identical to that of the standalone network (Methods). Figure 3b-d illustrate the “virtual” compressed size during the training process for the three texts. The compressed data sizes calculated based on this loss were 2.122 MB, 2.425 MB and 5.498 MB for the three texts, whereas the bzip2 compressed sizes were 2.660 MB, 2.808MB and 2.886 MB, respectively. For wiki0 and web, the compression efficiency of the composite network is separately 20.24% and 13.65% (Figure 3e).

Noting that Reformer is trained with texts from Wikipedia, the similarity between the PLM training set and wiki0 is much higher than that between the PLM training set and web. The similarity degree is regarded as the main reason for the difference of compression efficiency of the composite network. Consistently, pattern extraction for por using the composite network failed due to the scant similarity between Wikipedia text (English) and Portuguese (Figure 3e).

**Figure 3.**
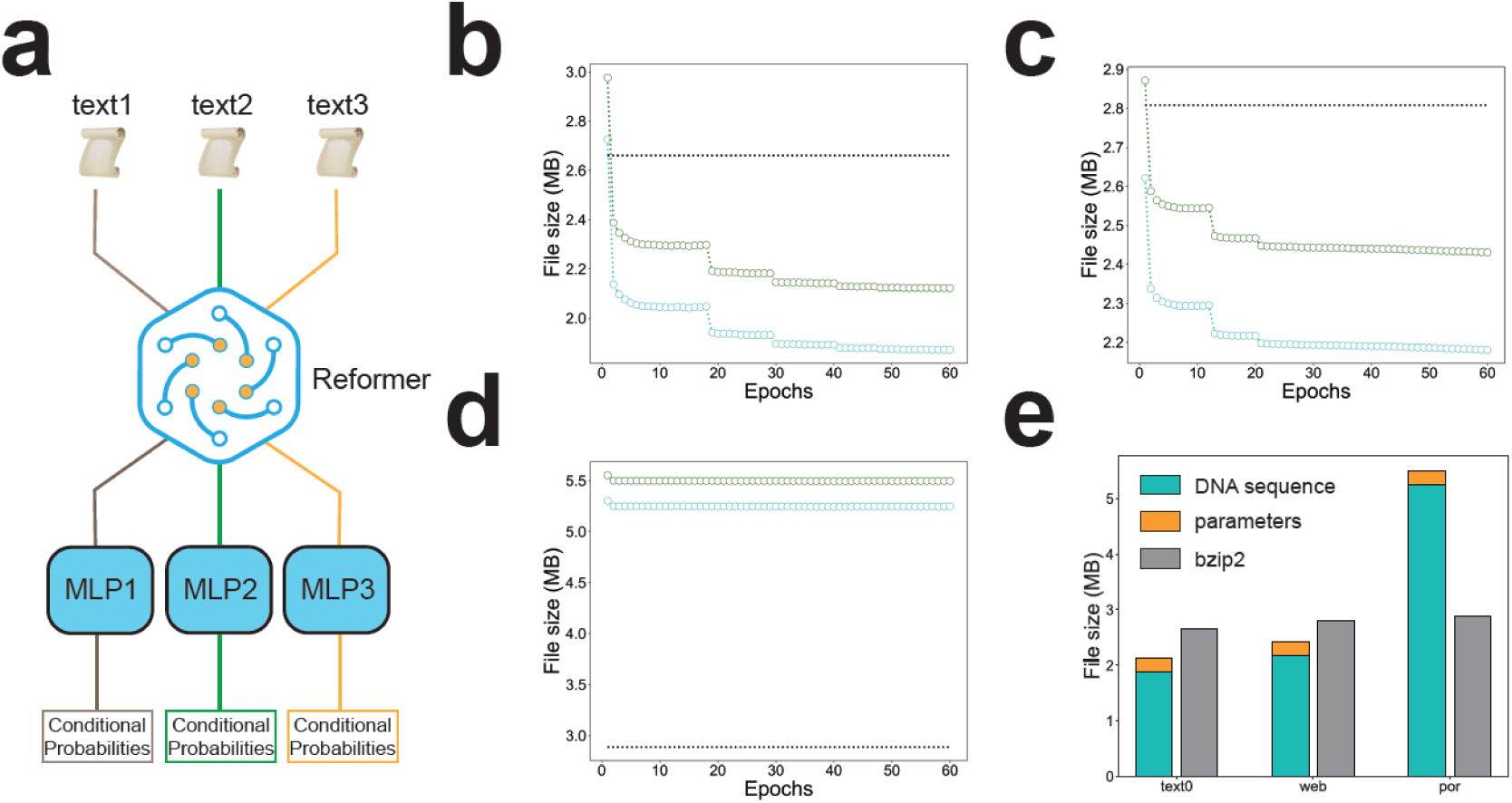
Pattern extraction facilitated by PLM. (a) Architecture of the composite network. (b)-(d) The “virtual” compressed size during the training process of (b) wiki0, (c) web and (d) por. In each panel, the cyan dots indicate the file size without the size of parameters, while the green dots indicate the file size with the size of parameters, and the black dash line indicates the file size compressed by bzip2. (f) The total compressed size of three texts, in comparison with bzip2.

### Base shifter for sequence transformation

After training of the networks, the original data were converted to a series of conditional probabilities. We designed a specialized module, called base shifter, to convert these conditional probabilities into nucleotide sequences. This module is comprised of two components, a probabilistic sequence register (PSR) and a nucleotide sequence converter (NSC). The PSR converted a string of conditional probabilities into a subset of the probability space, and then gradually recorded the corresponding nucleotide. The NSC was a hierarchical finite state machine that transformed the symbolic representation of the probability space in a bit-wise manner according to its state.

In the PSR, we developed base-4 arithmetic coding to adapt to the nucleotide space, which was a variant of conventional base-2 arithmetic coding [39–40]. One-dimensional probability space was used when we compressed one-dimensional sequences (*e.g.*, text). In this case, the space occupied by four nucleotides is a non-overlapping one-dimensional line, which together form the probability space (Figure 4a). The space occupied by each nucleotide was further divided to form higher-order representations of the probability space, where the corresponding spaces of the representations are adjacent but not coincident (Methods).

Along with the recording of nucleotides in the PSR, the NSC operated these nucleotides in a special Galois field GF{A, C, G, T} according to the current state of the NSC, resulting in a different representation. The NSC was a hierarchical finite state machine (FSM), which was comprised of two layers of FSMs. The bottom-layer FSMs defined the computing rules between two nucleotides. Each of the bottom-layer FSMs was composed of 4 nodes, and each node contained four edges that directed to another node (including itself). When a nucleotide inputted into a bottom-layer FSM, a state conversion was induced, and a nucleotide was simultaneously outputted. Figure 4b shows the structure of the “base” bottom-layer FSM, whose computing rules are illustrated in the bottom table. In this bottom-layer FSM, the output nucleotide was identical to the input nucleotide, regardless of the current state. However, in other bottom-layer FSMs, the output nucleotide was determined by both the input nucleotide AND its current state (Figure 4c). In top-layer FSM, each state was a bottom-layer FSM. The initial state of the top-layer FSM was the “base” bottom-layer FSM. However, when the input nucleotide was about to generate a sequence constraint, a state conversion was induced in top-layer FSM, leading to a change in the computing rules between nucleotides. Therefore, the input nucleotide was likely to generate a different output nucleotide, thereby avoiding the constraint (Figure 4d).

The path of state conversion in top-layer FSM was defined by a random number sequence, which was generated by a specific random seed. Recording the specific states after each conversion were unnecessary since the whole conversion path can be reproduced by the random seed. However, the positions of state conversion were indispensable for re-transformation to the original sequence. Therefore, the extra redundancy added during sequence transformation was depended on the times of state conversion, which was supposed to be minimized by an optimal random seed (supplementary note 3, Figure S2 and S3). As an illustration, we performed *in silico* simulation using base shifter, and the setting of constraints were same as that in DNA fountain [8]. Here we generated 10,000 random DNA sequences with a length of 256 bp (512 bits data with a coding density of 2 bit/base), and sent these sequences to base shifter for sequence transformation. For each pair of DNA sequence and random number, the size of redundancy required for de-transformation (*i.e.*, the random seed and the positions of state conversion) was recorded, where 256 random seeds were applied for each DNA sequence. Figure 4e shows the number of constraints and the number of state conversion for each pseudo-sequence. Noting that only the minimum value was substantial for each DNA sequence, since we can just take the random number of this value for sequence transformation. The average number of state conversion is ∼1 for each pseudo-sequence, which corresponds to a redundancy of 8 bits (*log*_2_256). The redundancy for random seed is also 8 bits (*log*_2_256). Therefore, only a redundancy ratio of (8+8)/(512+8+8)=3.03% was required to completely exclude the constraints in DNA sequence. For comparison, in DNA fountain, 4 out of 38 bytes (10.53%) was used to exclude these constraints for each DNA oligo [8].

**Figure 4.**
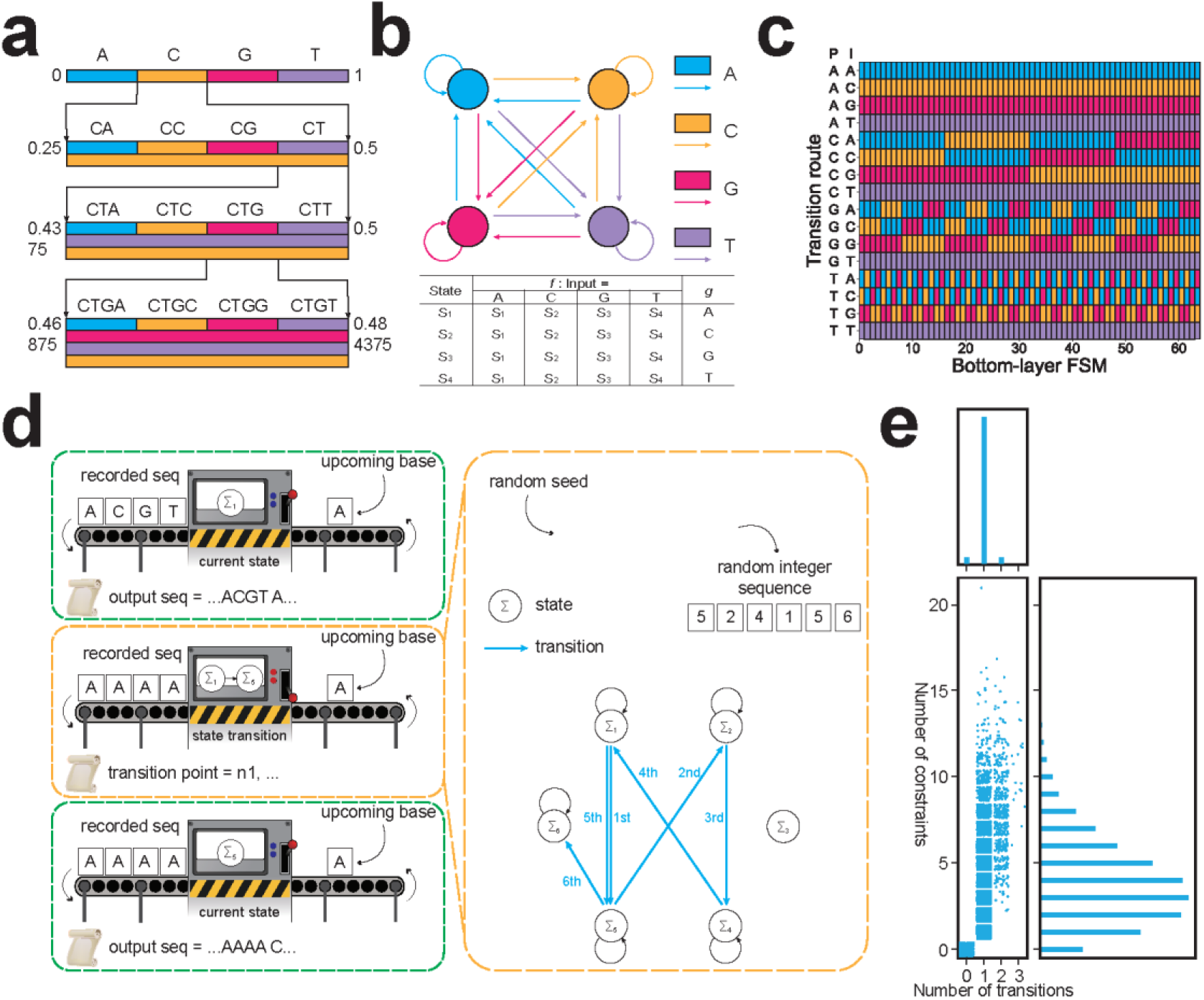
Design and performance of base shifter. (a) The mapping between probability space and nucleotide representation. (b) The computing graph and state transition table of “base” bottom-layer FSM. (c) The computing rules of the first 64 bottom-layer FSM. (d) The workflow of top-layer FSM. (e) Simulation results of 10,000 random DNA sequences with a length of 256 nt. For each sequence, 256 random seeds are used for simulation. Dots are randomly fluctuated to show density.

### Coding pipeline comparison and practical data storage

To better illustrate the superiority of FECDO, we encoded the three texts we used into nucleotide sequences. Meanwhile, we built a “conventional” coding pipeline, which was comprised of bzip2 for compression and DNA fountain for transcoding (constraint exclusion). Noting that we did not include error correction here, since it is independent from compression and transcoding. Given the pattern extraction results we have obtained, we used the composite network and base shifter for wiki0 and web, while used the standalone network and base shifter for por. After training, the parameters of the networks were stored, as part of the coding results. The text data were re-inputted into the network to obtain conditional probabilities for each position in the data sequence. Next, these probabilities were transformed into constraint-free DNA sequences with base shifter (Methods).

The original sizes of the three texts are all 10 MB. For the conventional coding pipeline, the length of DNA sequences was 11,892,632 nt for wiki0, 12,554,128 nt for web and 12,900,540 nt for por. In comparison, the length of DNA sequences coded by FECDO was 8,744,042 nt for wiki0, 9,993,375 nt for web and 11,292,105 nt for por. Remarkably, the improvement of compression ratio was 26.48% for wiki0, 20.40% for web and 12.47% for por. This result had verified the practicality of FECDO, namely, the potent pattern extraction by neural networks and efficacious constraint exclusion by base shifter.

To confirm the feasibility of FECDO in the whole cycle of DNA data storage (compression, encoding, DNA synthesis, storage, DNA sequencing, decoding, de-compression), we went through the whole process with data coded using FECDO. Here we chose the web text as the data for storage, whose initial size was 10 MB and encoded into 9,993,375 nt DNA sequences. To cope with the potential errors in synthesis and sequencing, we added error correction codes, which was a variant of MEPCAL that we previously developed [14], where we added appropriate amount of redundancy to cope with an error rate of 0.5% (including substitutions, insertions and deletions). The DNA fragments used for storage was 200 nt in length, which was comprised of primer region, bumper region, barcode region and payload region (Methods). The bumper region was added at the end of the fragments, since we found error hotspots in nanopore sequencing [14]. A total of 80,000 DNA fragments were used to carry all the information required for data retrieval, which were stored in *in vitro* DNA pools.

As for data retrieval, the DNA fragments in DNA pools were amplified by PCR, and then used for library preparation, and finally sequenced using MinION (Oxford Nanopore Technologies). After base-calling and reads clustering, the nucleotide sequences of reads were used for decoding and de-compression (Methods). The errors in nanopore sequencing were well resolved by MEPCAL decoder, to obtain error-free DNA sequences. Actually, we found that DNA sequences can be faithfully retrieved at even lower sequencing coverage, indicating a further decrease in sequencing cost.

The decoded DNA sequences, along with the random number and the positions for state conversion, were sent back to base shifter to restore the original DNA sequence. This sequence was then mapped to the probability space represented by nucleotides. The characters in the text were then iteratively restored. Here we preserved the first 100 characters as the first input of the prompt network with frozen weights, which led to a conditional probability of the 101^st^ position. By comparing this conditional probability and the sequence-mapped probability space, the character of the 101^st^ position was recognized, followed by the restoration of the 102^nd^ position and so on in a similar manner. At last, all data were restored without any error.

## Discussion

In this study, we described a delicate coding method for DNA data storage, FECDO. Neural networks were applied to effectively extract the inner pattern of digital data, and used for prediction during both encoding and decoding. Benefit from the mechanism of finite state machine and random seed, base shifter was devised to exclude arbitrary sequence constraints with minimal overhead. Combining these strategies, more efficient and flexible data coding in DNA was realized. The superiority of FECDO was demonstrated by separately coding real data using both FECDO and methods used in previous studies of DNA data storage. Notably, FECDO achieved better performance of compression, with a considerable decrease of 12-26% in the length of coded nucleotide sequences. This decrease was directly corresponded to the reduction of the cost of DNA synthesis, which has remained a main defect in recent years despite the development of related fields. As a source coding method, FECDO can be flexibly integrated with general-purpose or specialized error correction algorithm to achieve robust data storage in DNA. As an illustration, we integrated FECDO with our previously proposed DNA channel encoding algorithm MEPCAL, to encode a real text into DNA sequences. These sequences were synthesized and then sequenced with nanopore sequencing. The stored data was perfectly retrieved after error correction with a sequencing coverage of only ∼10×. The original texts were then restored by de-compression, and no error was observed.

Previous algorithms developed for DNA data storage mainly focused on error correction (nucleotide mutations and dropout of oligos) and sequence conversion (from digital stream to nucleotide sequences, with unsuitable sub-sequences excluded) [5–15]. While various types of digital data had been stored in DNA, these data were processed by traditional compression algorithms adapted from information science [18–20]. However, these methods often failed to realize “tight” compression towards the real information entropy [21–24]. For DNA data storage, this compression ratio is directly related to the expenditure, since the storage medium (*i.e.*, the DNA strands) must be *de novo* synthesized. Recently, a few studies tried to develop compression methods for DNA data storage, yet little substantial progress was made [41]. In contrast, FECDO exhibited excellent compression capability, as illustrated by the compression of real data.

For pattern extraction, we used two different approaches, namely, standalone network and pre-trained language models with auxiliary network. The performance of the former was slightly better than the commonly used compression method (bzip2), while the latter exhibited remarkable better compression efficacy for certain data. However, this efficacy relied on the similarity between the data to be stored and the training set of PLMs. In contrast, data compression using standalone network is more stable regardless of the contents of data. To fully utilize standalone network to compress idiographic or uncommon data, the architecture of the network is supposed to be optimized, to better extract the data pattern. For example, 2D CNN was supposed to be assembled into the network to extract the local features in images.

Sequence screening is essential for DNA data storage, for specific DNA sequences can cause severe damage to DNA synthesis and sequencing, thereby harming the reliability of data storage. However, sequence screening usually reduces the available coding space, thus reducing the storage density. For instance, Church *et al.* avoided adjacent identical nucleotides by mapping only 1 bit to 1 nucleotide [5]. This design resulted in a contraction of 50% of the coding space. Similarly, Grass *et al.* used RS codes based on GF(47), where three consecutive identical nucleotides were excluded [7]. The coding space was thereby contracted by 15%. Erlich *et al.* excluded specific subsequences by performing XOR with a stream of random numbers [8]. In this case, the coding space was complete, but a series of random seeds occupied a part of the payload region on the DNA oligos (10.53%). By using hierarchical finite state machine and random seed, we accomplished efficacious sequence screening with a minimum cost (∼3%), without directly contracting the coding space. As a result, we achieved higher storage density compared with previous DNA coding methods.

Retrospecting the development of DNA data storage in the past decade, the cost and speed have been remaining as the crux for both academic research and industrial production. While the writing and retrieving speed can be highly promoted by parallelism and novel technologies, the decrease in the cost was confined by the fundamental DNA synthesis process. Given this premise, data coding served as an essential procedure for reducing the cost. How to compress data into DNA with maximum efficiency, while guaranteeing exclusion of constraints and robust sequence recovery, remained to be further excavated.

